# Autosomal dominant CDC45 deficiency with allelic expression bias causes a novel genetic disease of the immune system

**DOI:** 10.64898/2026.03.04.709645

**Authors:** Nicole C. Guilz, Yong-Oon Ahn, Seungmae Seo, Madrikha D. Saturne, Matilde I Conte, Sana R. Shehzad, Everardo Hegewisch-Solloa, Luis Alberto Pedroza, Micah Castillo, Preethi Gunaratne, Ivan K. Chinn, James R. Lupski, Emily M. Mace

**Author notes:** Department of Natural Sciences, University of Maryland Eastern Shore, Princess Anne MD USA. Department of Pediatrics, Children’s Hospital of Philadelphia, Philadelphia PA USA.

## Abstract

Here we describe a damaging heterozygous variant in CDC45 in an individual with common variable immunodeficiency (CVID) and recurrent viral infections. This individual has variably decreased number of circulating NK cells, disruption in the ratio of CD56^bright^ to CD56^dim^ cells, and consistently decreased NK cell function. Interestingly, the inherited CDC45 variant is also present in a sibling with less severe clinical manifestations; we determined that allelic bias of the damaging allele accounts for this differential expressivity. As previously reported for other helicase variants that cause inborn errors of immunity (IEI), we found cell cycle defects in immune cells from the proband that lead to reduced survival of NK cells. Together, these findings link another member of the core replicative helicase complex to inborn errors of immunity and highlight the sensitivity of NK cells to these variants. They also define another case of allelic bias contributing to variable expressivity of an IEI gene.

## Introduction

Natural killer cell deficiency (NKD) is a primary immunodeficiency where natural killer cells are affected in the absence of other immune defects (1). Individuals with NKD present with malignancies and susceptibility to viral infection, especially from the Herpesviridae family (2). Though NKD is rare, four members of the CDC45-MCM-GINS (CMG) helicase are associated with NKD, including MCM4 (3, 4), GINS1 (5), MCM10 (6), and GINS4 (7, 8). In addition, other genes associated with cell cycle progression and DNA replication have been associated with inborn errors of immunity that include a component of NK cell dysfunction but may also have other immune or extra-immune dysregulation (9–11).

The CDC45-MCM-GINS (CMG) helicase is essential for eukaryotic replication and plays an important role in maintaining genomic stability (12). Cell division cycle protein 45 (CDC45) is a critical component of the CMG helicase that acts as a rate-limiting factor in DNA replication in mammalian cells (13, 14). As part of the helicase complex, CDC45 is responsible for initiation of DNA replication and DNA double helix unwinding (15, 16). Biallelic loss of CDC45 causes Meier-Gorlin syndrome (MGS7; MIM: 617063)17, a disorder characterized by short stature, small ears, and no or small patellae (17, 18). In addition to the role of CDC45 within the CMG helicase complex, CDC45 can also protect the genome by binding to single-stranded DNA (19) and DNA junction sites (20). As such, CDC45 plays a critical role not just in replication initiation, but in DNA unwinding and synthesis (16).

The unusual susceptibility of NK cells to damaging variants in the CMG helicase has been previously documented (3– 8). While the underlying cause of NK cell sensitivity to mild replication stress is not completely understood, it is likely related to differences in the expression of CMG complex members and differences in response to replication stress between NK cells and T cells. NK cells express lower levels of CMG helicase proteins upon entering cell cycle and have a lower threshold for apoptosis in the presence of replication stress than T cells (21). Commitment to the NK cell lineage is associated with cell proliferation, and in CMG helicase patient cells this leads to loss of NK cell progenitors and decreased NK cells in peripheral blood (8). As CMG genes are required for embryonic viability and more severe variants frequently cause Meier-Gorlin Syndrome, it is unlikely that variants causing NK cell deficiency are demonstrating an NK cell-specific function of this complex (22). Instead, the degree of loss of function appears to lead to degrees of immune involvement, and this degree of loss of function can be further complexified by the presence of allelic bias (8). While incompletely understood, autosomal random monoallelic expression, or allelic bias, has recently been shown to contribute to variable penetrance in inborn errors of immunity (23). This includes CMG helicase variants, namely GINS4, where allelic bias has been shown to lead to variable disease penetrance in two siblings with inherited compound het-erozygous variants (8). Here, we describe a case of complex immunodeficiency with a significant NK cell component arising from loss of a single allele of CDC45 and compounded by allelic bias.

## Results

### Clinical phenotype and immune features

The female proband (P1, Fig. 1A) presented with recurrent episodes of sinusitis, pneumonia, and Giardia in adolescence and was diagnosed with common variable immunodeficiency (CVID) at the age of 18. She has consistently near-absent IgA and IgM (Supp. Table 3) and remains on bimonthly intravenous immunoglobulin treatment. While she is CMV- and EBV-negative, the proband has had recurrent episodes of disseminated varicella zoster (VZV) with post-herpetic femoral neuropathy, which have required hospitalization and antiviral prophylaxis. She has also had treatment resistant bacterial infections including an aggressive pseudomonal otitis and chronic mastoiditis complicated by postauricular fistula, neck abscess, left-sided deafness and the need for temporal dural repair, requiring prolonged courses of intravenous antibiotics. She has had multiple episodes of meningitis with lymphocytic pleocytosis and colonization with enteropathogenic E. coli. She has significant lymphoid hyperplasia throughout the gastrointestinal tract, including an obstructing lesion which required a right hemicolectomy with resection of ileocecal valve and resulted in vitamin B12 deficiency and malabsorption. Flow cytometry on the lesion showed predominantly T lymphocytes, fewer B lymphocytes and 1% NK cells (CD16+CD56+). She also has biopsy-proven cirrhosis due to primary biliary cholangitis (AMA-negative). Her cirrhosis has remained well-compensated since diagnosis. SARS-CoV-2 infection required hospitalization with sepsis, acute hypoxic respiratory failure and cardiac complications but was cleared at day 83 after prolonged antiviral treatment. Other viral infections of note include human metapneumovirus and norovirus. Following the onset of the COVID-19 pandemic in 2020, the proband has maintained social distancing and essentially been homebound. Clinical immunophenotyping shows chronic lymphopenia with consistently low CD4+ T cells, occasionally low B cells, and low NK cell numbers in the peripheral blood (Table 1 and Supp. Table 4). Mitogen responses and antigen stimulation were largely found to be normal apart from low responses to tetanus antigen (Supp. Table 5).

**Table 1.** Selected longitudinal laboratory values. For data from additional timepoints see Supp. Table 2. L, low; H, high.

|  | Reference Range (X10<sup>3</sup>/UL) | 1990 | 1991 | 2013 | 2016 | 2017 |
| --- | --- | --- | --- | --- | --- | --- |
| WBC | 4.5 – 11.0 |  |  |  |  |  |
| ALC |  |  |  |  | 1,176 | 663 |
| CD3+ | 798 – 2,594 | 604 | 1174 | 931 | 953 | 537 (L) |
| CD3+CD4+ | 579 – 1,841 | 394 | 699 | 533 | 497 (L) | 280 (L) |
| CD3+CD8+ | 184 – 855 | 234 | 412 | 329 | 300 | 212 |
| CD19+ | 63 – 461 | 129 | 309 | 87 | 94 | 46 (L) |
| CD20+ | 59 – 457 | 131 | 344 | 86 | 82 | 46 (L) |
| CD3-CD56+CD16+ | 89 – 472 |  |  |  | 118 | 80 (L) |
| CD3+CD56+CD16+ | 0 – 200 |  |  |  | 165 | 106 |
| CD2+ | 884 – 2,800 | 723 (L) | 1240 |  | 1,047 | 590 (L) |
| CD56+ | 137 – 478 | 50 (L) | 64 (L) | 43 (L) | 306 | 186 |
| CD8+CD56+ | 30 – 200 |  |  |  | 223 (H) | 133 |

**Fig. 1.**
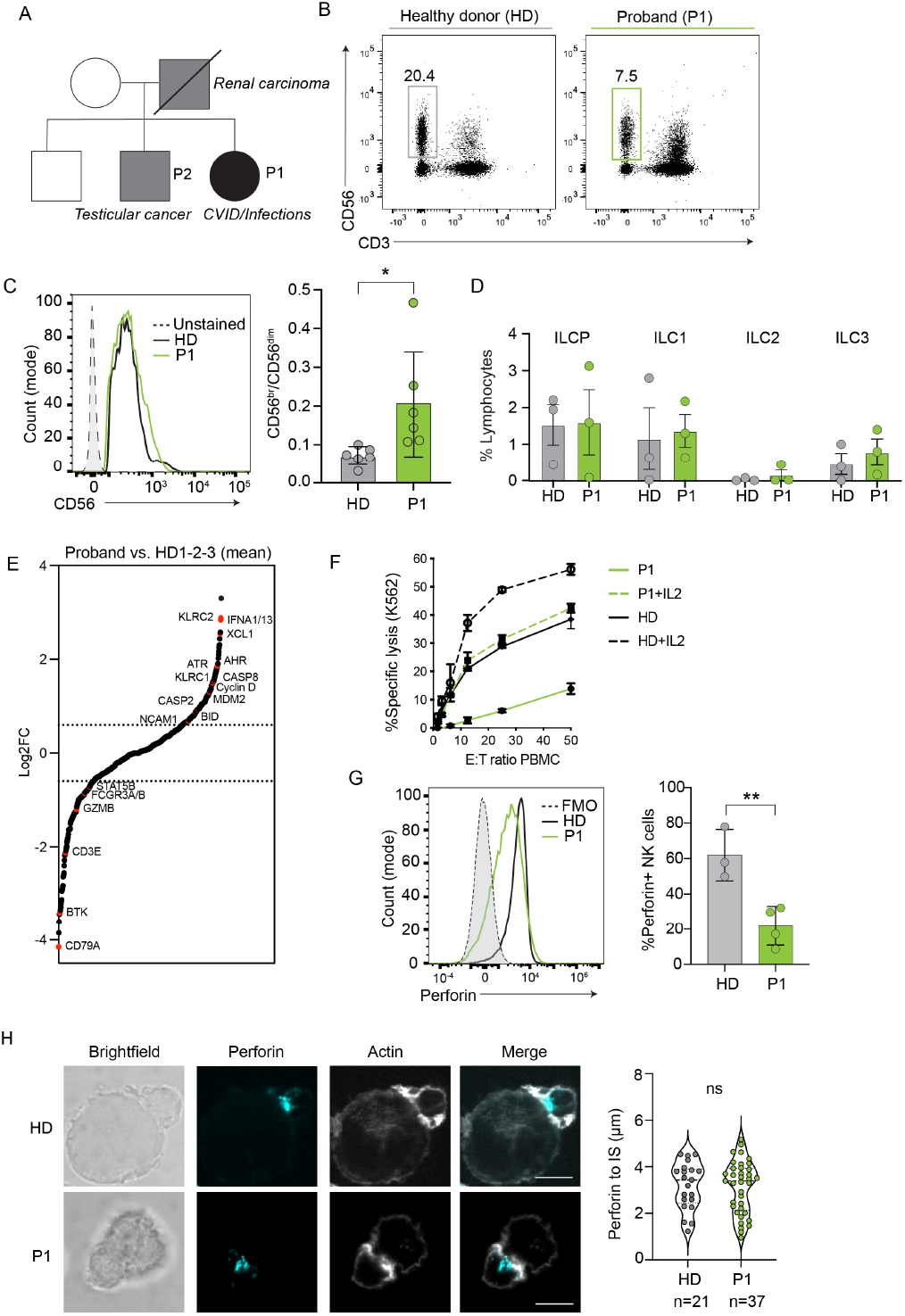
Impaired NK cell function and abnormal NK cell phenotype. An adult woman with CVID and NK cell deficiency was repeatedly evaluated for NK cell function and phenotype. A) Pedigree of the proband (P1) and her family. B) Representative flow plots of NK cells from peripheral blood gated on CD56+CD3-NK cells. C) Representative CD56 histogram (left) and quantification of the ratio of CD56^bright^ to CD56^dim^ cells (right); meanSD of 6 technical replicates and 6 biological replicates (healthy donors). *p<0.05 by Student’s unpaired two-tailed t-test. D) Frequencies of circulating ILC subsets and ILC progenitors from P1 (3 technical replicates) and healthy donors (3 biological replicates). E) NK cells were enriched from peripheral blood of the proband (P1) and an age-matched healthy donor (HD) and gene expression was measured by Nanostring (see Supp. Table 4 for list of genes). Heatmaps represent relative expression of 27 genes selected from those >1.5-fold different between proband and healthy donor. F) NK cell cytotoxicity against K562 target cells of proband (P1) and healthy donor (HD) in the absence and presence of IL-2. Shown is one representative assay performed in triplicate, mean±SEM. G) Perforin in CD56+CD3 NK cells measured by flow cytometry; histogram is one representative from 4 independent sampling times for the proband quantified on the right. mean±SD, *p<0.05 by unpaired two-tailed Student’s t-test, 3 biological replicates for healthy donors. H) Representative confocal microscopy images of primary NK cells conjugated to K562 target cells; actin (gray) and perforin (cyan). Distance of lytic granules to the immune synapse; ns, not significant by unpaired Student’s two-tailed t-test; n=21 (healthy donor, HD) or n=37 (proband, P1).

The proband’s family history is significant for her father having metastatic transitional cell carcinoma of the renal pelvis and her brother (P2) having a polycellular testicular cancer cured with surgery (Fig. 1A). In addition, P2 experienced recurrent multiple colonic adenomatous polyposis since 2003 at the age of 46, with a lifetime total >86 polyps. Genetic testing in 2011 for APC variants and in 2017 for APC and MUTYH variants were negative for pathogenic mutations or variants of uncertain significance. No gene deletions or duplications were detected. Given the significance of the father’s history, repeat genetic testing was subsequently performed to expand beyond the adenomatous polyposis panel to include a renal/urinary tract cancers panel. A likely pathogenic variant was identified in FH. While heterozygous for autosomal recessive fumarate hydratase deficiency, inherited heterozygous variants in FH have been reported and are associated with familial cancers, including renal cell carcinoma (24). This individual also experiences chronic otitis media with petrous and mastoid air cell opacifications and layering fluid on the left side without associated bone destruction. He had an unremarkable SARS-CoV-2 infection but reports frequent oral herpesviral lesions. Notably, this individual is also EBV- and CMV-negative.

### NK cell phenotype

In addition to the involvement of other immune lineages, including CD20+ B cells and CD4+ T cells, an impairment of natural killer cells was suspected due to low NK cell numbers in peripheral blood of the proband on repeated clinical evaluation and her recurrent and severe herpesviral infections. NK cell phenotyping and functional analysis at timepoints from 2016 to 2022 (ages 46-55 years) showed a variable frequency of NK cells in peripheral blood that were lower than healthy donors but not absent (Fig. 1B). While NK cell frequencies and numbers were variable, an imbalanced ratio of increased CD56^bright^ to CD56^dim^ NK cells was also noted (Fig. 1C). Despite these differences, we detected no abnormalities in the low frequencies of other circulating innate lymphoid cell subsets found in peripheral blood (Fig. 1D).

To better define the NK cell phenotype in the proband, NK cells were enriched from peripheral blood for Nanostring gene expression analysis of 595 immune-related genes (Supp. Table 6). Comparison of the proband’s NK cells to NK cells from three unrelated healthy donors identified 152 genes differentially expressed >2-fold, including those associated with NK cell development and function (NCAM1, CD3E, GZMB, FCGR3A/B, KLRC1, KLRC2, AHR, ZBTB16), cytokines and cytokine signaling (STAT5B, IFNA1/13, XCL1), and cell cycle and apoptosis (BID, ATR, CASP8, Cyclins D and E, MDM2) (Fig. 1E, Supp. Table 7).

We tested the functionality of the proband’s NK cells and found that cytotoxicity against K562 target cells was decreased compared to controls but could be increased in response to IL-2 (Fig. 1F). We also found decreased frequencies of perforin-positive NK cells and decreased perforin mean fluorescence intensity (MFI) in longitudinal measurements of proband NK cells (Fig. 1G). This was additionally validated by the decrease in PRF1 (-1.15-fold) and GZMB (-2.29-fold) gene expression in proband NK cells relative to healthy donor cells detected by Nanostring gene expression analysis (Fig 1E).

Effective NK cell cytotoxicity is dependent on formation and function of the immunological synapse32. To test whether the decreased NK cell function from proband NK cells arose from impaired immune synapse formation, we performed confocal microscopy of NK cells and K562 target cells. NK cells formed conjugates with target cells and, while perforin levels were qualitatively decreased in NK cells from the proband, quantitative analysis revealed no significant differences in the average distance between perforin-containing granules and the immune synapse (IS) (Fig. 1H). This suggests that the critical steps of immune synapse formation by proband NK cells are intact despite target cell killing on a population level being impaired.

### Phenotype of peripheral blood cells defined by single cell sequencing

To further define the peripheral blood phenotype of the proband, we performed scRNA-Seq on freshly isolated PBMCs from the proband (P1), her brother (P2), and an unrelated healthy donor control. Transcripts from individual cells were barcoded via the 10x Genomics Chromium platform and sequenced for 5 gene-expression profiling. Following quality control validation, clustering was performed using Seurat (25). Cluster identities were manually assigned based on previously defined immune cell identities using the pooled sample as a reference (Fig. 2A-B, Supp. Fig. 2). Cluster 4 (NK cells) were found at a similar frequency in the proband sample and expression of genes defining the NK cell cluster was similar to that in the healthy control and P2 (Fig. 2C, D). The proband sample (P1) had distinguishing features that included the over-representation of activated lymphoid and myeloid populations (clusters 6 and 8; 32.6% and 20.8%) that were only represented at low frequencies (<1%) in other samples (Fig. 2C). We noted a decreased frequency of some lymphocyte subsets in the proband sample, including B cells and CD8 and CD4 T cells. However, a robust population of JUN, JUNB, JUND, CD69, and NFKBIA expressing, activated lymphoid cells (‘cluster 6’) was present in the proband sample. PANTHER pathway analysis identified activation and apoptosis signaling as hallmarks of this population (Fig. 2E). We identified a similar population of myeloid cells expressing genes associated with activation (‘cluster 8’, Fig. 2E). Both these populations had increased absolute numbers of cells and frequencies in the proband sample. In some cases, higher expression of genes associated with these clusters was also noted (Fig. 2F).

**Fig. 2.**
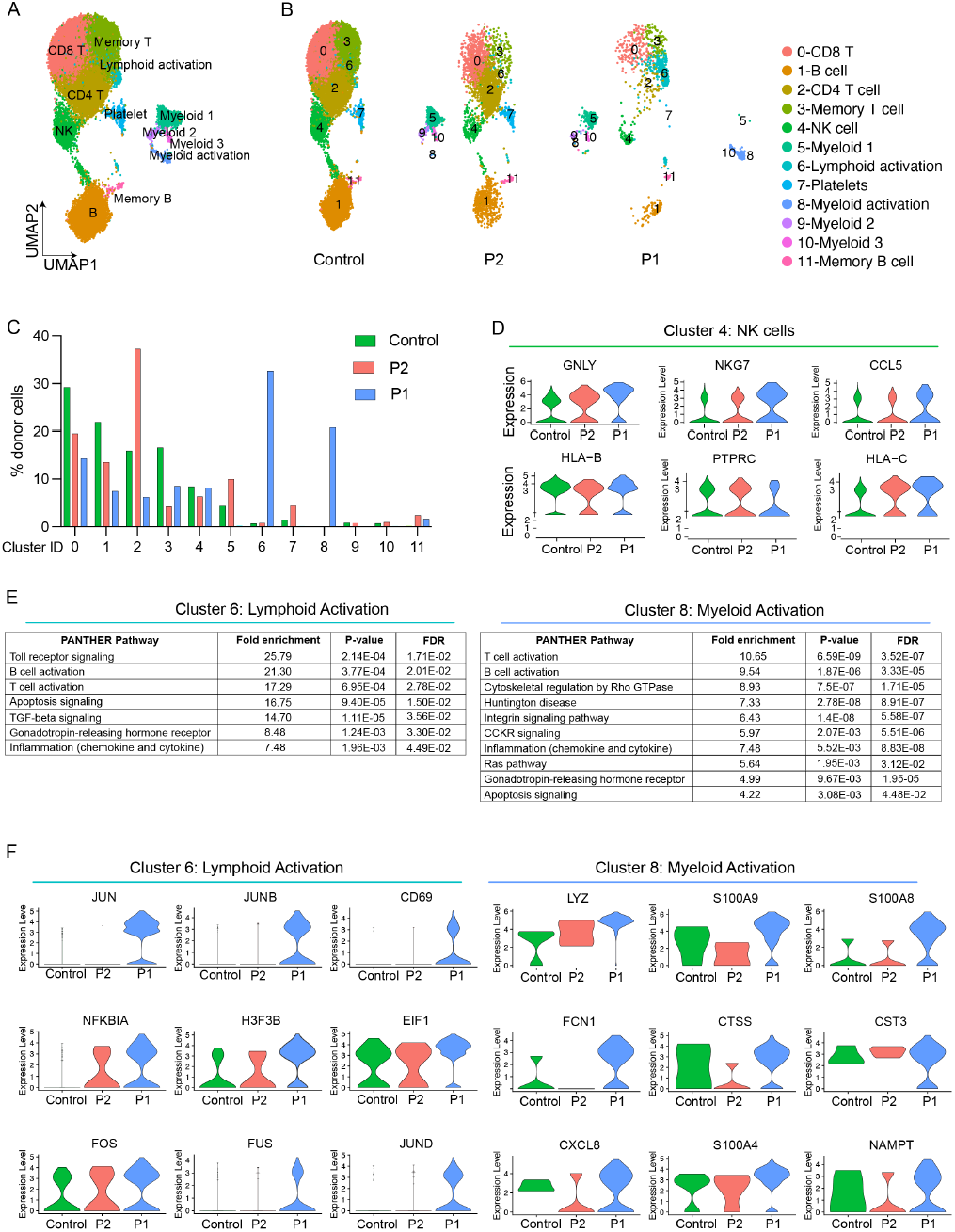
scRNA Seq demonstrates increased frequency of activated myeloid and lymphoid cells in proband peripheral blood cells. PBMCs from the proband (P1), her brother (P2) and an unrelated healthy donor were analyzed by scRNASeq (10X Genomics). Analysis was performed by Seurat v.3. and cluster identities were defined by differential gene expression (see also Supp. Fig. 2). A) UMAP and cell type assignment of all 44,367 cells from all donors. B) UMAP from (A) separated by donor. C) Frequency of each cluster shown colored by donor. D) Violin plots showing expression of top 6 expressed NK cell genes in cluster 4 from each donor. E) Pathway analysis of cluster 6 and cluster 8 performed by PANTHER analysis. FDR, false discovery rate. F) Expression of top 9 genes from clusters 6 and 8 split by donor.

### Identification and validation of a CDC45 variant leading to loss of protein expression

Exome sequencing (WES) was performed using DNA from blood samples of the proband (P1), her mother, and her sibling (P2). A heterozygous frameshift variant was found in exon 4 of CDC45 resulting from an insertion of two nucleotides (c.287_288insTG, p.F96fs). This result was confirmed in the proband with Sanger sequencing (Fig. 3A) and was also present in her brother (P2) but not the mother, suggesting it was inherited from the proband’s deceased father (Fig. 1A). This variant was not previously found in ClinVar, EXaC, or gnomAD and is predicted to result in nonsense-mediated decay (26–28). CDC45 binds directly to the MCM2-7 hexamer and the GINS tetramer to form the CMG complex (15). Together with recruitment of the GINS tetramer, CDC45 is required for transformation of the inactive MCM hexamer on double-stranded DNA to the active CMG complex (13, 15). As such, loss of its expression is expected to directly impact CMG function. To better understand the effect of this variant on protein expression, immortalized B cell lines were created from PBMCs from healthy donors and the two siblings carrying the CDC45 frameshift variant. Decreased expression of CDC45 protein was found in the proband cells compared to five healthy donors and no truncated CDC45 protein was detected (Fig. 3C, Supp. Fig. 3A). Western blotting of BLCLs from the proband’s brother with the CDC45 variant (P2) also demonstrated decreased detection of CDC45, confirming that this variant negatively affects CDC45 protein expression (Fig. 3C). In contrast with P1, NK cell phenotype and function from P2 were within normal range upon repeated measurements of this individual’s cells (Supp. Fig. 3B-D). This was consistent with a milder clinical phenotype that was limited to recurrent oral herpesviral infections.

**Fig. 3.**
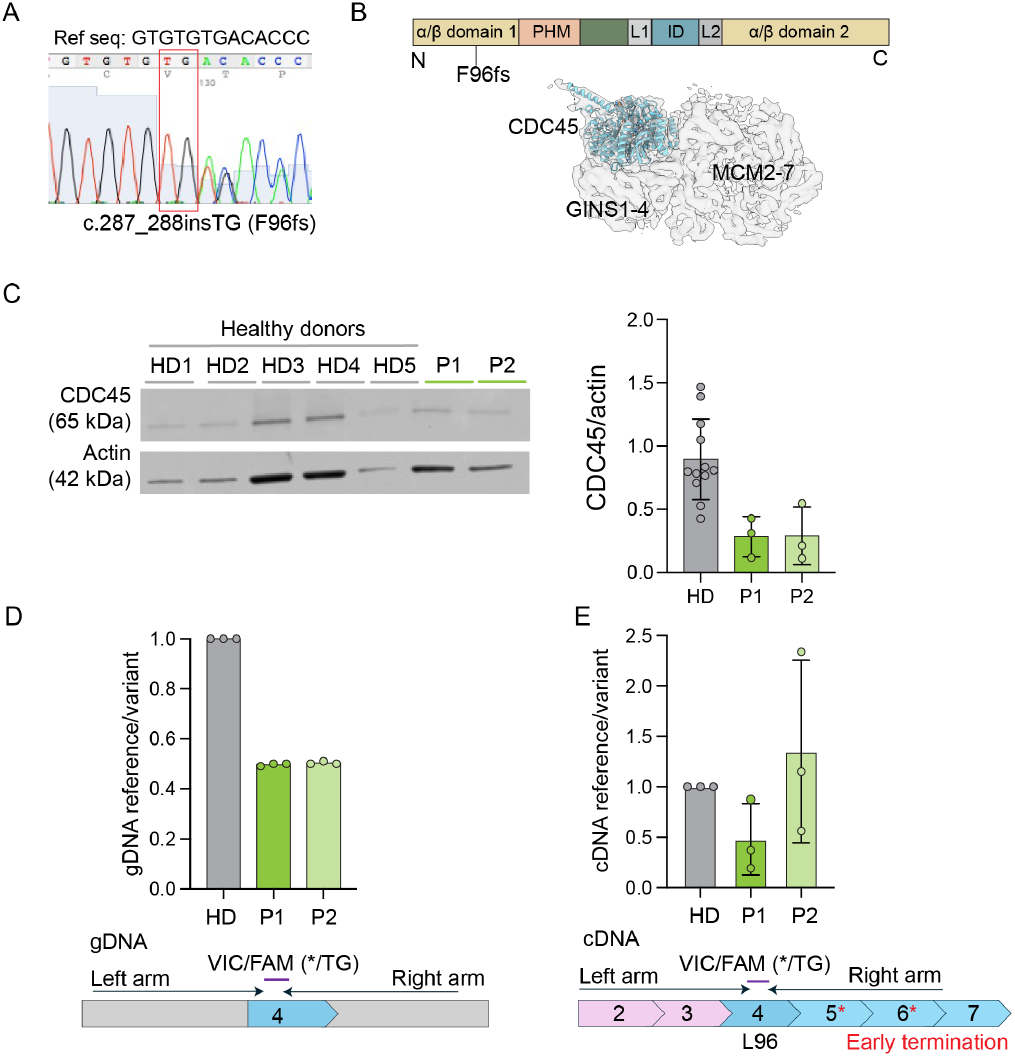
CDC45 variant identified by whole exome sequencing. A) Sanger trace of DNA from the proband showing TG nucleotide insertion and frameshift at c.287 in CDC45. B) Structure of CDC45 showing location of the F96fs variant. PHM, protruding helical motif; L1, linker 1; ID, all-helical interdomain; L2, linker 2. Placement of CDC45 within the CMG protein complex relative to MCM2-7 visualized with ChimeraX also shown. C) Western blotting of 5 healthy donor (HD1-5), P1 and P2 B cell lines (BLCLs) probing for CDC45 and actin. CDC45 intensity relative to actin is quantified on the right; mean±SD, 3 technical replicates per donor. D) Detection of the CDC45 reference allele detected using probes for genomic DNA from an unrelated healthy donor (HD), P2, and P1. Data from 3 technical replicates are normalized to HD. E) Detection of the reference variant using probes for mRNA performed in parallel with the genomic detection in (D) from HD, P2, and P1; shown are 3 independent replicates normalized to HD, mean±SD.

Given the observation that P2 had a milder clinical phenotype, and given the recent description of allelic bias as a contributing factor to variable expressivity of IEI genes (8, 23), we asked whether allelic bias could contribute to differences between the proband and her sibling with the same CDC45 genotype but less severe clinical phenotype. Using probes for genomic DNA (gDNA), we measured the ratio of reference to variant allele and confirmed 50% expression of the CDC45 reference allele in P1 and P2 BLCLs, whereas an unrelated healthy donor had 100% expression of the reference allele, as expected (Fig. 3D). Using the same samples to measure cDNA, we found higher expression of the reference allele cDNA in the sample from P2 relative to the proband (Fig. 3E). While these amounts were variable, the consistently increased expression of the reference allele in P2 cells suggests that allelic bias could contribute to the variable penetrance of disease in this family.

### CDC45 deficiency leads to impaired cell cycle progression and increased DNA damage

CDC45 is a core member of the CMG complex that regulates DNA replication (29). We first considered the cell cycle profile of BLCLs derived from the CDC45-deficient proband and found increased frequency of cells in S phase compared to the mean of 10 healthy donor lines (Fig 4A). Cells from P2 had a frequency of cells in S phase consistent with healthy donor lines (Fig. 4A, B). Expression of MCM6, a replisome protein that is up-regulated during S phase, in total lysate from cells from the proband was not different relative to 5 healthy donors and P2, further validating that cells were activated and entering S phase (Fig. 4C).

**Fig. 4.**
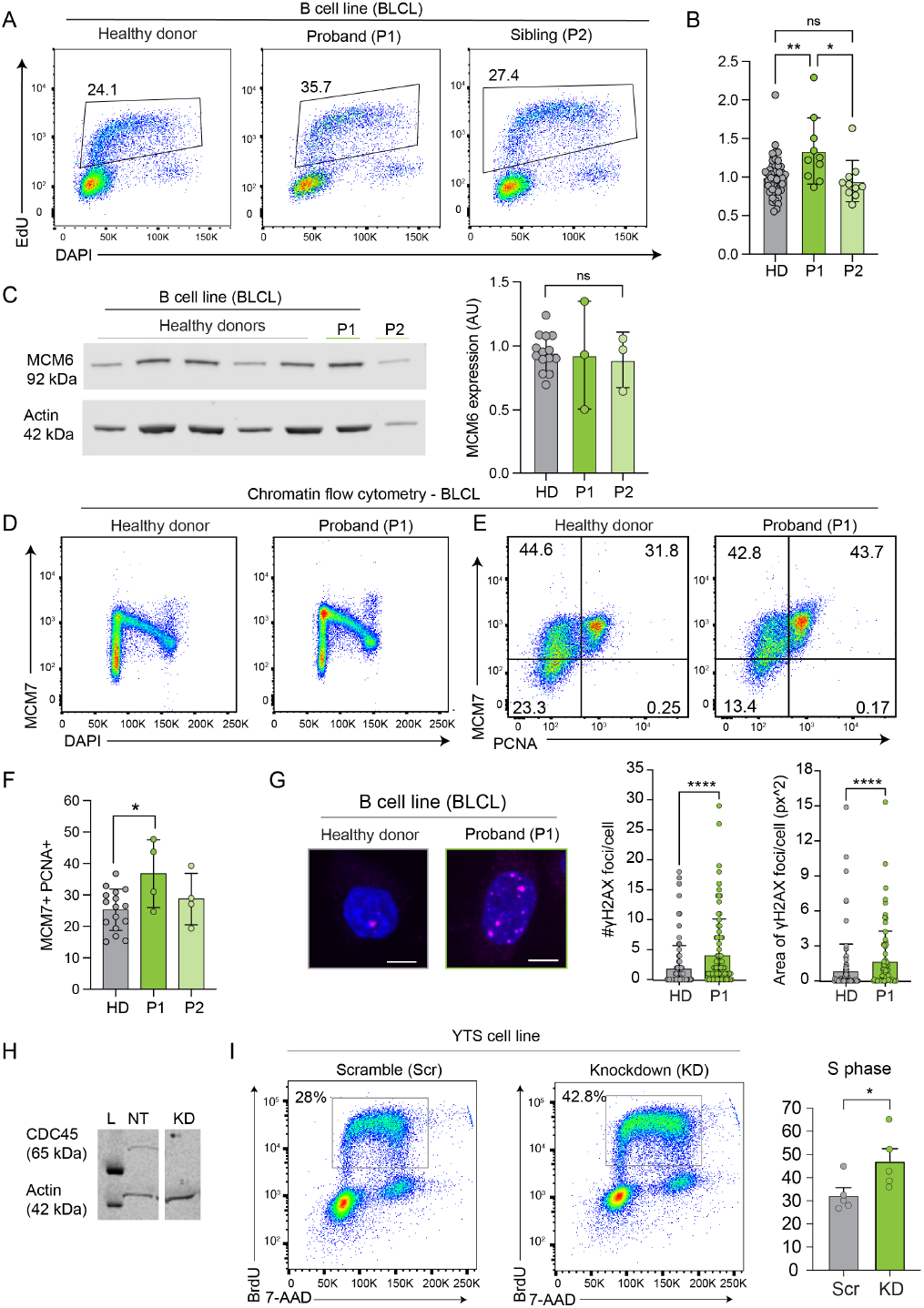
Impaired cell cycle progression and increased replication stress in CDC45-deficient cells. A) Representative cell cycle analysis of immortalized B cells (BLCL) from an unrelated healthy donor, proband (P1) and the sibling (P2) measured by flow cytometry. B) Quantification of the frequency of cells in S phase relative to the mean of HD for each independent replicate. Ten technical replicates are shown for P1 and P2, n=47 from 10 biological replicates for HD. mean±SD, ordinary one-way ANOVA with multiple comparisons. C) MCM6 protein expression measured in 5 healthy donors, P1 and P2. Representative Western blots and quantification of the ratio of MCM6/actin from 3 technical replicates (P1 and P2) is shown. N=14 for HD (4-5 biological replicates performed as 3 technical replicates to correspond with P1 and P2 replicates). mean±SD, not significant by ordinary one-way ANOVA. D) Chromatin flow cytometry showing loading of MCM7 (G1) and unloading of MCM7 as DNA is replicated as detected by DAPI staining (S phase). E) Chromatin flow cytometry showing chromatin-bound MCM7 and PCNA from healthy donor and proband (P1) samples. F) Quantification of MCM7+ PCNA+ cells from 4 technical replicates for HD, P1, and P2; n=16 HD samples (4 biological replicates), mean±SD, p=0.0406 by one-way ANOVA with multiple comparisons. Significant individual comparisons are marked. G) γH2AX in BLCLs imaged by confocal microscopy (left) and quantified from 3 technical replicates (right). n=87 (healthy donor), 101 (proband); mean±SD, ****p<0.0001 by unpaired Student’s t-test. Scale bar = 5 um. H) Validation of shRNA-mediated knockdown of CDC45 in the YTS human NK cell line with non-targeting (scramble, Scr) as a control. Representative of 3 technical replicates. I) Representative cell cycle analysis of CDC45 knockdown (KD) or non-targeting control (Scr) YTS cell lines. H) Quantification of the frequency of cells in S phase measured by flow cytometry as in (I). n=5 technical replicates, *p<0.05 by Student’s unpaired two-tailed t-test.

DNA replication begins with assembly and activation of the CMG complex, followed by fork firing and elongation. These events can be monitored by measuring the association of the replisome with chromatin as an indication of binding to DNA prior to activation (30). To test whether CDC45 deficiency affects the loading of the replisome onto chromatin, we performed chromatin flow cytometry to measure binding of MCM7 to DNA prior to fork firing (30). We observed no qualitative difference in the ability of cells from the proband or her sibling (P2) to load MCM7 onto chromatin (Fig. 4D). This observation was further supported by quantifying co-expression of chromatin-loaded MCM7 and PCNA as a read-out for origin licensing, which defined a higher frequency of PCNA+ MCM7+ on chromatin of P1 cells (Fig. 4E, F). To-gether with the increased frequency of cells found in S phase in the proband-derived lines, this suggested that CDC45 is required for fork firing and elongation, but not expression of other helicase components or replisome assembly.

To determine whether the presence of this variant led to induction of DNA damage responses, we performed confocal microscopy of 2AX using BLCL lines in the absence of additional replication stress. Microscopy and quantitative analysis identified a significantly increased frequency and area of H2AX foci in cells from the proband relative to the healthy donor control lines (Fig. 4G). Finally, we generated a CDC45 knockdown NK cell line to test the effect of decreased CDC45 expression on cell cycle progression. Lentiviral shRNA guides targeting CDC45 or with a non-targeting sequence were stably transfected into the YTS cell line and knockdown was confirmed by Western blot (Fig. 4H). Consistent with our data from BLCL lines, we found a significantly increased frequency of cells in S phase in the knockdown condition relative to the non-targeting control (Fig. 4I).

### Effect of CDC45 variant on NK cell proliferation

Given the impaired cell cycle progression and NK cell phenotype in the proband, we tested NK cell proliferation in response to homeostatic cytokines. We cultured proband or healthy donor PBMCs in IL-15 to induce NK cell proliferation. After 3 days of IL-15 culture, we found that proband NK cells robustly proliferated, as did healthy donor controls (Fig 5A). While NK cells had only undergone 2-3 rounds of division at 3 days, it appeared that a greater frequency of proband cells had entered cell division relative to healthy donor cells (Fig. 5A). We then isolated primary NK cells and repeated IL-15 stimulation, this time for 5 days to better capture multiple rounds of cell division (Fig. 5B). Again, we found that proband NK cells robustly proliferated and entered cell cycle. While we did not detect significantly increased apoptosis in proband NK cells in response to IL-15 alone, we next asked whether mimicking NK cell viral activation would adversely affect NK cell survival. As NK cells activated with cytokine-induced memory-like conditions undergo apoptosis in response to mild replication stress (21), we performed CIML activation of primary NK cells from proband or healthy donors. Proband NK cells in CIML cytokines had decreased viability relative to healthy donors, while IL-15 stimulation alone led to a notable but not statistically significant decrease in proband cells (Fig 5C). Together, these data suggested that activation with CIML cytokines, which mirrors viral activation, led to decreased NK cell survival in the proband. While difficult to definitively show, this could account for the variably decreased NK cell number in the peripheral blood of the proband.

**Fig. 5.**
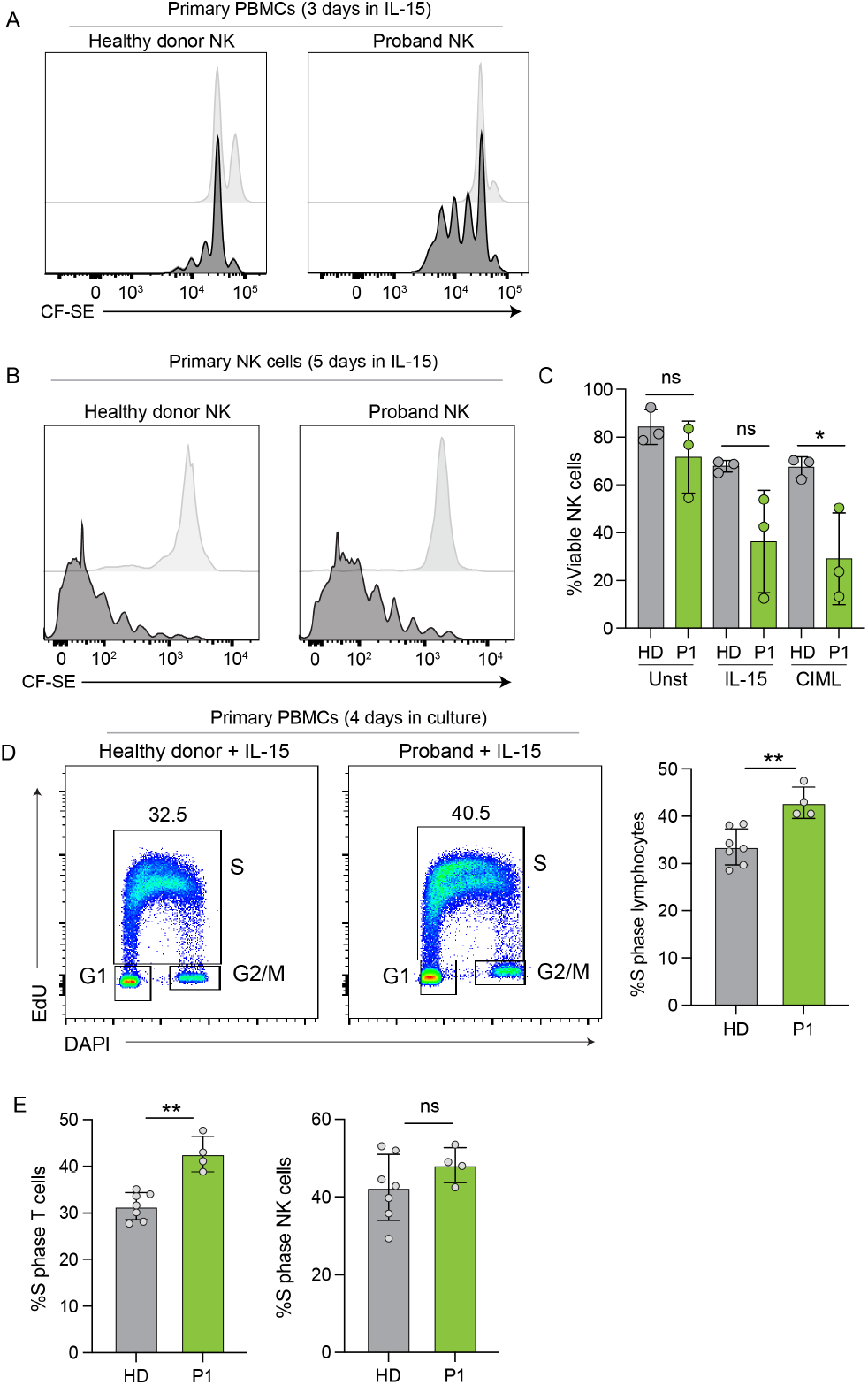
CDC45-deficient primary NK cells have decreased viability in response to activation. Primary PBMCs or NK cells from healthy donor or P1 were labeled with CF-SE and cultured with 5 ng/ml IL-15 for 3-5 days. A) Representative histograms showing cell divisions of primary PBMCs. B) Representative histograms showing cell divisions of primary NK cells. C) NK cells were activated with cytokine-induced memory-like (CIML) NK cell conditions or IL-15. Cell viability after 5 days measured by flow cytometry using viability dye, mean±SD. D) Representative cell cycle plots of all lymphocytes from PBMCs activated for 3 days with 5 ng/ml IL-15; 4 technical and biological replicates (healthy donors) are quantified to the right, mean±SD, **p<0.05 by Student’s unpaired two-tailed t-test. E) Quantification of cells in S phase gated on CD3+ T cells or CD56+CD3-NK cells. 4 technical and biological replicates (healthy donors) are quantified to the right, mean±SD, **p<0.05 or not significant (ns) by Student’s unpaired two-tailed t-test.

Finally, to better link cell proliferation with cell cycle entry, we repeated IL-15 stimulation of PBMCs. Cell cycle analysis of peripheral blood mononuclear cells stimulated with IL-15 for 4 days showed a higher frequency of lymphocytes in S phase in the proband sample than in healthy donor samples (Fig. 5D). When gating on NK cells and T cells, we found that the frequency of proband cells in S phase was increased in both subsets, but this difference was only significant in T cells, not NK cells (Fig. 5E). Together, these data suggest that increased frequency of cells in S phase is not unique to NK cells, but that decreased viability of NK cells in response to cytokine activation can contribute to the NK cell phenotype found in the proband.

### Decreased efficiency of NK cell differentiation from proband-derived iPSCs

To more closely study the effect of CDC45 variants on NK cell differentiation, iPSCs were generated from proband and healthy donor PBMCs and validated for pluripotency and karyotype. iPSCs were then differentiated into NK cells using the spin-EB method (31) (Fig 6A). The first stage of NK cell differentiation from iPSCs is CD34+ hematopoietic progenitor generation, and we found a decreased frequency of CD34+ cells in the proband line relative to healthy donor control line (Fig. 6B). As previously described, unrelated healthy donor lines consistently generated populations that were >50% mature NK cells (Fig. 6C). In contrast, proband-derived iPSCs generated very few mature NK cells apart from a single experiment in which mature NK cells were produced (Fig. 6C). While this finding was surprising, it was reminiscent of previous studies of allelic bias that resulted in similar observations from GINS4-deficient iPSC-derived NK cells (8). Further characterization of the NK cells from the proband that were generated in this experiment demonstrated decreased granzyme B expression relative to those of the healthy donor control line (Fig. 6D). This finding was reminiscent of decreased expression of lytic granule components in primary NK cells from the proband.

**Fig. 6.**
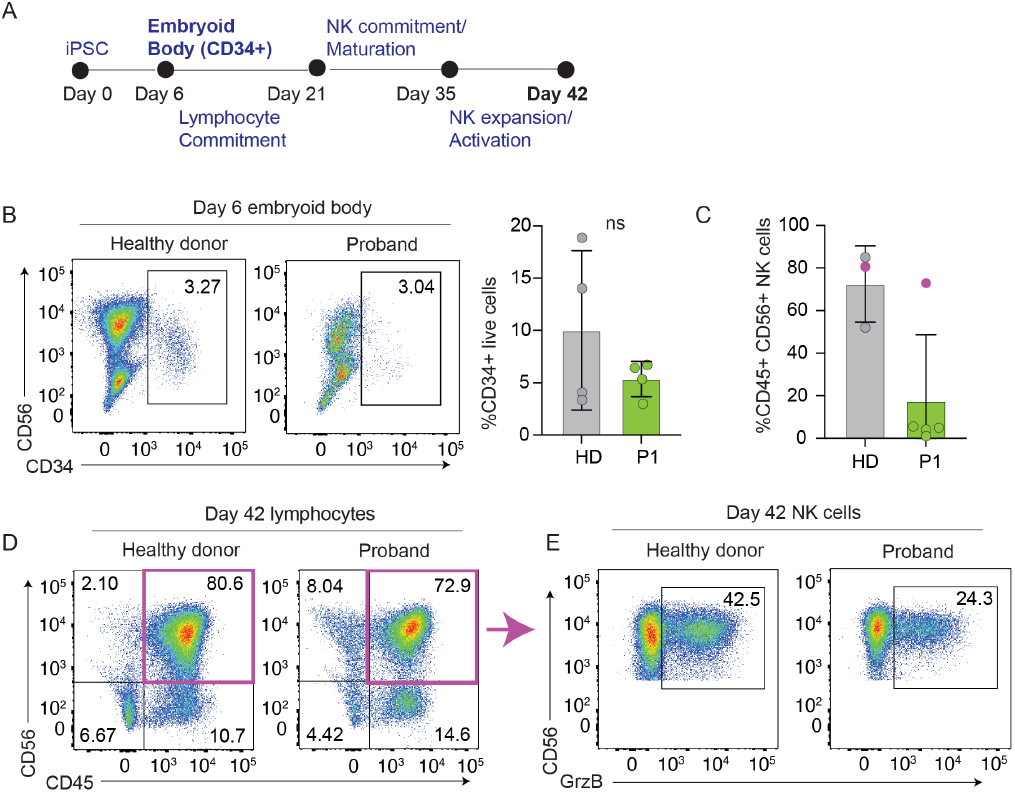
Decreased efficiency of NK cell differentiation from proband-derived iPSCs. Blood cells from the proband or an unrelated healthy donor were reprogrammed to iPSCs and validated. Differentiation to CD34+ hematopoietic progenitors then NK cells was performed using a feeder-free culture system. (A) Schematic of NK cell differentiation from iPSCs showing major stages of NK cell generation. Day 6 and day 42 time points were sampled for this study. B) Representative flow cytometry plots and frequency of non-mesoderm (CD56) CD34+ cells at day 6; gated on live cells. Quantification of 4 independent replicates shown to the right; ns, not significant. C) Frequency of CD45+CD56+ NK cells at day 42 from 4 independent technical replicates. Magenta data point indicates the experiment shown in (D). D) Flow cytometry plots of CD45+CD56+ NK cells at day 42; gated on live cells. E) Representative flow cytometry plots showing frequency of granzyme B+ (GzmB) NK cells at day 42. Gated on live cells and CD45+CD56+ cells.

## Discussion

Our description of an individual with a single damaging variant in CDC45 expands the phenotype of variants in the eukaryotic DNA helicase that are known to cause human NK cell deficiency. It also ascribes an immune deficiency to the effect of a single inherited pathogenic variant in CDC45. Biallelic CDC45 variants cause Meier-Gorlin Syndrome, including a distinctive phenotype with craniosynostosis and anorectal malformation in addition to the typical triad of short statue, microtia, and no or under-developed patellae (32, 33). No immune involvement has been reported to date in these patients. DiGeorge Syndrome, also known as 22q11.2 deletion syndrome, also often includes deletion of CDC45 (34, 35). Immunodeficiencies and autoimmune diseases have been reported in these patients, although other genes found on chromosome 22 likely contribute to this effect and make it difficult to isolate the effect of deletion of a single copy of CDC45 (35, 36). Our data suggests that loss of CDC45 expression can be associated with a clinical phenotype, although the allelic bias and resulting incomplete penetrance likely contribute to this outcome. As other immune cell lineages, including B cells and T cells, are affected in our proband, we consider this case of CDC45 deficiency to be an inborn error of immunity (IEI), but not exclusively an NK cell deficiency. T cells in the proband had increased length of S phase and impaired tetanus mitogen response, suggesting that they may also be affected in the proband. This observation based on clinical findings is supported by scRNA-Seq data showing decreased frequency of CD4+ T cells, CD8+ T cells, and B cells. Here, we focused on the NK cell phenotype to better understand the role of CMG complex members in NK cell biology that is highlighted by IEI. As in other cases describing sensitivity of NK cells to replication stress resulting from helicase variants (8, 21), we find that NK cells undergo apoptosis in response to cytokine-induced activation. We propose that the source of isolated NK cell deficiency in some individuals with helicase variants is a combined effect of the sensitivity of NK cells to mild replication stress and variants that affect NK cell homeostasis and function because they can be compensated for in other cell types by innately higher expression of the protein. The broader effect of CDC45 may reflect a lower tolerance for loss of its function, given its role as a rate-limiting component of the replisome13.

CDC45 is an essential gene, and its deletion leads to embryonic lethality (37). As demonstrated by the syndromic features of patients with Meier-Gorlin Syndrome, biallelic damaging variants lead to a range of developmental phenotypes. Evaluating this individual’s immune phenotype over time allowed us to identify variability in the numbers of NK cells in peripheral blood. Decreased frequency of NK cells was accompanied by overrepresentation of immature NK cells, suggesting impaired terminal maturation or homeostasis of mature NK cells. Differentiation of NK cells from iPSCs led to decreased frequency of mature NK cells. It also appeared to decrease the frequency of CD34+ progenitors, although variability in the healthy donor control prevented this difference from being statistically different. This is a different phenotype from that of GINS4 deficiency, in which the generation of CD34+ progenitors is not affected but impaired cell cycle progression leads to apoptosis of NK/innate lymphoid cell precursors (8). Instead, it is more similar to MCM10 deficiency, in which CD34+ progenitor differentiation is impaired from iPSCs with a haploinsufficient MCM10+/ phenotype (38).

The difference in clinical phenotypes between two siblings with the same genotype was initially difficult to reconcile. However, we found allelic bias that favored the variant allele in the proband, leading to less cDNA expression than predicted in the proband. This allelic bias conversely favored the unaffected allele in the proband’s sibling, leading to a greater than predicted protein expression in his cells. Whether this difference is the sole cause of the significantly more severe clinical manifestation in the proband is still unclear and difficult to determine. However, the existence of other rare instances of CDC45 single allele loss that have not been associated with NK cell deficiency, either in DiGeorge syndrome or because of other rare stop-gain mutations in CDC45 (26), suggests that loss of CDC45 expression on a single allele does not necessarily lead to haploinsufficiency.

While no other predicted pathogenic variants that segregated to result in disease were identified by next-generation sequencing, another variant not yet discovered cannot be ruled out. Further, we hypothesize that environmental factors, such as severe viral infections that the proband experienced in adolescence, could have contributed to the manifestation of disease in the proband. While the finding of allelic bias was surprising, recent studies have described monoallelic expression as a cause of incomplete penetrance in IEI (23). Notably, this includes in the case of GINS4 deficiency, where allelic bias in a family with biallelic variants contributes to a variable clinical and cellular NK cell phenotype between siblings (8). The mechanisms of allelic bias in the context of immunity and IEI remain incompletely understood but likely relate to features of lineage- and gene-specific properties.

In summary, the description of IEI linked to CDC45 provides another example of immunodeficiency linked to impaired replisome function and expands the effect of CDC45 insufficiency to include immune dysregulation. While the effect of CDC45 deficiency is broader than some previously described replisome proteins, such as GINS4 and MCM10, the sensitivity of this immunodeficiency to gene dose effects likely reflects sensitive thresholds of its tightly regulated expression during cell cycle progression. Our findings underscore the importance of understanding lymphocyte cell cycle dynamics and how these are differentially regulated between immune cell subsets.

## Methods

### Cell isolation and cell lines

Blood was collected under the guidance of the Institutional Review Board at Columbia University and in accordance with Declaration of Helsinki guidelines. PBMCs were isolated by density gradient centrifugation from peripheral blood of the proband or her sibling or the buffy coat of healthy donors from the New York Blood Center using Ficoll-Paque (Fisher Sci Cat 45-001-750). Buffy coats and healthy donors were and sex-matched when possible. PBMCs were stored frozen in 90% FBS 10% DMSO and thawed for each experiment. Once thawed, PBMCs were resuspended and maintained in RPMI 1640 medium (Thermo Cat 11875135) supplemented with 10% heat-inactivated human AB serum (GeminiBio Cat 100-512), 1% penicillin-streptomycin (Thermo Cat 15140163), 2 mM GlutaMAX (Thermo Cat 35050079), 1X non-essential amino acids solution (Thermo Cat 11140076), 1 mM sodium pyruvate (Thermo Cat 11360070), and 1M HEPES (Thermo Cat 15630130). YTS cell lines (RRID CVCL D324) were maintained with the same media as PBMCs, except with 10% heat-inactivated FBS (Atlanta Biologicals Cat S11150) instead of human serum. K562 cells (RRID CVCL 0004) were maintained in RPMI medium supplemented with 10% FBS, 1% PenStrep, and 2mM GlutaMAX. All cell lines were incubated at 37C with 5% CO2 and confirmed negative for My-coplasma prior to experiments.

### NK cell stimulation

Frozen PBMCs were thawed and briefly rested in human serum-supplemented RPMI medium. To activate NK cells using cytokine-induced memory-like (CIML) protocol (39), cytokines were added to the media for overnight stimulation (100 ng/mL IL-15, 50 ng/mL IL-18, 10 ng/mL IL-12; all cytokines from Peprotech). PBMCs were then washed three times and resuspended with fresh media and low-dose IL-2 (20 U/mL) for expansion phase in a round-bottom 96-well plate. For IL-15 stimulation, cells were cultured with 5 ng/ml IL-15 following labeling with CF-SE (Thermo Cat 34554) per the manufacturer’s protocol.

### Flow cytometry

For analysis of NK cell phenotype, including ILC subsets, flow cytometry panels are listed in Suppl. Table 1. Cells were washed in PBS then surface stained with fluorochrome-conjugated antibodies and viability dye at 4C for 20 minutes while protected from light. Cells were washed again then fixed and permeabilized for intracellular staining using the FoxP3 nuclear staining kit (Fisher Sci Cat 00-5523-00). Fluorescence minus one and/or unstained controls were included for all experiments. Data were acquired on a BD Fortessa and analyzed with Flow Jo v.10.9.0 (BD Biosciences, RRID SCR 008520). Cell cycle analyses were conducted with a BrdU assay kit (BD Biosciences Cat 559619, RRID AB 2617060). Cells were incubated with 10 M BrdU for two hours followed by extracellular marker staining, fixation, denatured with DNase, and incubation with anti-BrdU. Cells were stained with 7-AAD for five minutes, then analyzed on a BD Fortessa cytometer (RRID SCR 018655).

Chromatin-bound proteins were detected by chromatin-flow cytometry (40) with minor modifications. Briefly, BLCL lines were pulsed with 10 µM EdU for 30 min, then harvested, washed, and resuspended with RNase extraction buffer on ice for 10 min. Cells were washed, fixed, and stained with antibodies against MCM7 (RD Systems, clone 2068B, IC9217G), PCNA (BioLegend Cat 307908, RRID AB 314698), and Histone H3 (BioLegend Cat 641012, RRID AB 2565890). EdU incorporation was detected by click reaction (Thermo Fisher Cat 10643, Click-iT Plus Alexa Fluor 647 Picolyl Azide Toolkit). After intracellular staining, cells were cultured with Alexa Fluor 647-picolyl azide (2 uM), CuSO4 (100 mM), L-ascorbic acid (20 mg/ml) in PBS for 30 min. After wash, cell suspension was stored in flow cytometry buffer with DAPI (3 uM) and RNase A (25 ug/ml) overnight and analyzed using BD LSR Fortessa X-20 (RRID SCR 019600).

### Digital PCR

Alle-specific gene expression was measured using QuantStudio Absolute Q Digital PCR System (Thermo) according to the manufacturer’s instruction. Custom TaqMan SNP genotyping probes for the CDC45 variant and reference were designed based on a target nucleotide (reference-VIC/variant-FAM) with 200 base pairs of left and right arms (probe sequences listed in Supp. Table 2).

To validate allele-specific gene expression, cDNA gBlocks gene fragments of CDC45 for the reference allele or the variant were synthesized (Integrated DNA Technologies) and tested as positive controls (Supp. Fig. 1). gDNA from BLCLs was isolated using Quick-DNA Microprep Kit (Zymo Research) and 80 ng of gDNA was used in the dPCR reaction. Total RNA was isolated using Quick-RNA Microprep Kit (Zymo Research) cDNA was further generated from the DNase I-treated RNA using LunaScript RT SuperMix (New England Biolabs). cDNA from 250, 50, and 10 ng of total RNA were used for dPCR to determine optimal lambda values. 10 µl of PCR mixture was prepared with gDNA or cDNA, 5x PCR master mix, and 40x probe/primer mix (TaqMan Gene Expression Assay). 9 µl of PCR mixture loaded into the plate and covered with 15 µl of isolation buffer and gaskets, and reference/variant (VIC/FAM) quantification was determined by cp/µl. All samples passed QC with ROX signal.

### Confocal microscopy

NK cells were briefly co-cultured with K562 target cells for 15 to 45 minutes in an eppendorf tube, then gently transferred to a poly-L-lysine coated microscope slide. Cells were fixed and permeabilized using Cytofix/Cytoperm (BD Biosciences Cat 554714) and then immunostained with phalloidin Alexa Fluor 568 (Thermo, Cat A12380) for actin and anti-perforin antibody (BioLegend Cat 308110, RRID AB 493254).

Cells were adhered to 1.5 imaging chambers followed by fixation and permeabilization using Cytofix/Cytoperm (BD Biosciences Cat 554714). Immunostaining was performed with anti-γH2AX Alexa Fluor 647 (BD Biosciences Cat 560447, RRID AB 1645414). DAPI staining was performed for 10 minutes prior to imaging. Images were acquired with a 100X 1.46 NA objective on a Zeiss AxioObserver Z1 microscope stand equipped with a Yokogawa W1 spinning disk by imaging cell volumes with a 0.17 µm Z step size. Illumination was by 405 nm, 488 nm, and 647 nm solid state lasers and detection was by a Prime 95B sCMOS camera. Data were acquired in SlideBook software (Version 6, Intelligent Imaging Innovations) and exported as OME-TIFF files for further analysis. Images were analyzed as Z projections in Fiji (41) (RRID SCR 002285) using the “Analyze Particles” plug-in with a minimum size of 0.05 µm^2^. Data were exported to Prism 10 (GraphPad Software, RRID SCR 002798) for graphing.

### NK cell cytotoxicity assays

Cytotoxicity of PBMCs against K562 erythroleukemia target cell line was performed by four-hour Cr51 release assay. Briefly target cells were incubated with Cr51 radionuclide (Perkin Elmer, NEZ030S001MC) for one hour, washed and incubated with PBMCs at increasing effector to target ratios, or in the absence of effectors for spontaneous release controls, in triplicate at 37C for 4 hours. After 4 hours, total experimental release controls were lysed with 1% octylphenoxy-polyethoxyethanol. Plates were spun and the supernatant was transferred to a LumaPlate (PerkinElmer), dried overnight, and read in a gamma scintillation counter. Total lysis and spontaneous lysis controls were included. Percent specific lysis was calculated using the following formula: [(experimental release-spontaneous release)/(total releasespontaneous release)] X 100.

### Western blots

Cells were lysed in RIPA buffer (Thermo Cat 89900) with 1X Halt Protease and Phosphatase Inhibitor cocktail (Thermo Cat 78443). Cell lysates were separated by gel electrophoresis using a 4%-12% gradient cell (Thermo Cat NW04120BOX) or 12% gel (Thermo Cat NW00127BOX) and then transferred to a 0.2 pore-size nitrocellulose membrane (Thermo Fisher Cat LC2000). The membranes were blocked using 5% nonfat dry milk solution (Fisher Scientific Cat 50-488-786) in PBS-tween 0.1%, then incubated with primary antibodies (CDC45 D7G6, Cell Signaling Technology Cat 11881, RRIDAB 2715569; beta-actin, Sigma-Aldrich Cat A5441, RRID AB 476744; MCM6 Abcam Cat ab201683, RRID AB 2924827) in a 1% BSA or nonfat dry milk solution. Secondary antibodies were applied in a 1% nonfat dry milk solution (IRDye 800CW Goat Anti-Rabbit IgG LiCOR Cat 926-32211, IRDye 680RD Goat Anti-Mouse IgG LiCOR Cat 926-68070, IRDye 800CW Goat Anti-Mouse IgG LiCor Cat 926-32210, IRDye 680RD Goat Anti-Rabbit IgG LiCor Cat 926-68071). Membranes were imaged on the LiCOR CxL and analyzed using the ImageStudioLite software (version 5.2.5).

### Generation of NK cells from iPSCs

Human iPSCs were generated from PBMCs from the proband or unrelated adult healthy donor controls using StemRNA Gen Reprogramming Kit (Stemgent)6. Cells were validated for pluripotency and periodically karyotyped and tested for mycoplasma. NK cell differentiation from iPSC was performed using the spin-embryoid body (EB) method25,29. Approximately 9000 iPSCs were plated in 96 well polystyrene round-bottomed microwell plates for EB formation. Cells were cultured in APEL-2 media (StemCell Technologies, Cat 5275) containing 10µM ROCK inhibitor (Y27632, StemCell Technologies, Cat 72302), 40ng/ml SCF (Peprotech, 300-07), 20ng/ml BMP (RD, 314-BP), 20ng/ml VEGF (RD, 293-VE), and 1µM GSK inhibitor (CHIR99021; StemCell Technologies, 72052). For spin-EB differentiation, at day 6 4 EBs were transferred to one well of a gelatin-coated 24 well plate with NK cell differentiation basal media supplemented with 5 ng/ml IL-3 (1st week only; Peprotech, Cat 200-03), 10 ng/ml IL-15 (Peprotech, Cat 200-15), 10ng/ml Flt3L (Peprotech, Cat 300-19), 20ng/ml SCF, and 20ng/ml IL-7 (Peprotech, Cat 200-07). NK cell differentiation basal media is made with 55% DMEM (Fisher Scientific, Cat 11965118), 28% Hams F12 (Fisher Scientific, Cat 10-080-CV), 15% human AB serum (GeminiBio, Cat 100-512), 1% L-Glutamax (Fisher Scientific, Cat 35050079), 1% Penicillin/streptomycin (100U/ml), 25µM BME (Fisher Scientific, Cat 21985023), 50 µM Ethanolamine (Sigma, Cat E0135-100ML), 25 µg/mL Ascorbic acid (Sigma, Cat A5960-25G), and 5 ng/mL sodium selenite (Sigma, Cat S5261-25G). After transfer to gelatin-coated plates at day 6, EBs were incubated for 35 days with weekly media changes and flow cytometry to monitor NK cell maturation by flow cytometry.

### Nanostring transcriptomic analysis

NK cells were isolated from whole blood of the proband and 3 unrelated healthy donors and were submitted for Nanostring multiplex gene expression analysis of about 600 human immune-related genes (nCounter Human Immunology V2). Total RNA was isolated using manufacturer’s instructions (Macherey-Nagel Nucleospin RNA XS Cat NC0389511). Raw gene expression values were exported from nSolver Analysis Software v.4.0 and the fold change of the proband values relative to the mean of all 3 healthy donors was calculated and converted to Log2FC for graphing with Prism 10 (GraphPad software).

### scRNA-Seq and analysis

PBMCs were thawed in a water bath at 37°C and washed with warm culture medium. Thawed cells were then spun at 500 rpm for five minutes, washed once, spun at 500 rpm for five minutes once more, then re-suspended in PBS (w/o calcium and magnesium) with 0.04% BSA. Washed cell suspensions were loaded onto a 10X Genomics chip N, following the Chromium Next GEM Single Cell 5’ HT Reagent Kits v2 (Dual Index) protocol (CG000423|Rev C). Modular kits for the 10X Chromium Connect were used to automate library preparation from cDNA. Gene Expression (GEX) sequencing libraries were generated using the Chromium Next GEM Automated Single Cell 5’ Reagent Kits v2 user guide (CG000384|Rev D). Libraries were assessed for quality using a High-Sensitivity D5000 chip on an Agilent 4200 TapeStation and were quantified with a Qubit Flex Fluorometer (Thermo Fisher Scientific). Libraries were sequenced on a NovaSeq 6000 (Illumina, RRID SCR 016387) to obtain 150 base paired end reads. The 10X Cloud Analysis portal was used to run cellranger v7.0.0 on all FASTQ files. Sequencing data was aligned to the GRCh38 human reference genome. UMAP dimension reduction, clustering, and cell cycle profiling was done using Seurat (v4.3.0.1, RRID SCR 016341)30.

### Statistics

Statistical analyses were conducted using Prism 10 (GraphPad, RRID SCR 002798). All data show mean±SD except where noted. One-sample 2-tailed Student’s t-test was used to compare the mean of experimental conditions to a control. One-way ANOVA was used to compare multiple conditions. P values on graphs are represented as: *p<0.05, **p<0.01, ***p<0.001, ****p<0.0001.

## ACKNOWLEDGEMENTS

We would like to thank Dr. Jordan Orange for sharing reagents and providing critical feedback on this project. We would also like to thank the staff of the Columbia Stem Cell Initiative Flow Cytometry Core Facility, under the leadership of Michael Kissner, at Columbia University Irving Medical Center for their contributions to the work presented in this manuscript. These studies also used the resources of the Herbert Irving Comprehensive Cancer Center Flow Cytometry Shared Resources funded in part through Center Grant P30CA013696. This work was supported by R01AI189818 to EMM, UM1HG006542 grant to JRL, and the Pate Foundation to EMM and NCG. YOA was supported by a Walkers Fellowship through the Edward P. Evans Center for Myelodysplastic Syndromes at Columbia University. We extend our deepest gratitude to the proband and her family for participation in this study.

## Notes

### Competing Interest Statement

The authors have declared no competing interest.

