## Supplemental Material for "Autosomal dominant CDC45 deficiency with allelic expression bias causes a novel genetic disease of the immune system"

**Supplemental figure 1. Generation and validation of allele-specific probes for genomic DNA (gDNA) and cDNA.** A) gBlock gene fragments were synthesized and used to validate allele-specific probes for the reference allele or F96fs allele. Sequences for gDNA or cDNA are shown as indicated. B) Synthesized gBlocks were used to test specificity of digital PCR probes. Probes were tested with WT gBlock or F96fs gBlock as indicated. Dot plots show output from digital PCR. C) Probes were tested with cDNA generated from BLCLs as indicated. Lack of amplification of the F96fs sequence likely reflects nonsense-mediated decay of residual CDC45 gene product.

**Supplemental figure 2. Unsupervised clustering of scRNASeq to assign cluster identities.** All donors were combined to generate the UMAP shown in Fig. 2A. Cluster identities were manually defined using expression of cluster-specific genes.

**Supplemental figure 3.** A) Uncropped Western blot showing CDC45 and actin as a loading control (correlated to Fig. 3C). B) Primary PBMCs from P2 or an unrelated healthy donor (HD) were incubated with K562 target cells in the presence or absence of IL-2 as indicated. Release of Cr51 was used to calculate specific lysis of target cells. A reference range of specific lysis values from >200 healthy donors is also shown (grey shaded region). C) Flow cytometry phenotyping shows expected frequencies of CD56+CD3- NK cells in peripheral blood. D) Frequency of perforin+ NK cells from unrelated healhy donors (HD), the proband (P1), and the proband’s sibling (P2). E) Frequency of NK/ILC precursors (Lin-CD117+), immature NK cells (CD56bright CD3-) and mature (CD56dimCD3-) cells from healthy donor (3 biological replicates), P2 (two independent evaluations and P1 (4 independent evaluations).

**Supplemental Table 1.** Antibodies used for flow cytometry panels

**Supplemental Table 2.** gBlock fragments used to synthesize probes for digital PCR

**Supplemental Table 3.** Longitudinal immunoglobulin levels from the proband.

**Supplemental Table 4.** Longitudinal

**Supplemental Table 5.** Mitogen and antigen responses from proband and control

**Supplemental Table 6.** Gene list from nCounter Human Immuno panel used for Nanostring analyses.

**Supplemental Table 7.** Differentially expressed genes (>2-fold) in the proband relative to mean of 3 healthy unrelated donors.
